# KmerKeys: a web resource for searching indexed genome assemblies and variants

**DOI:** 10.1101/2021.05.17.444256

**Authors:** Dmitri S. Pavlichin, HoJoon Lee, Stephanie U. Greer, Susan M. Grimes, Tsachy Weissman, Hanlee P. Ji

**Author notes:** Co-first authors (Dmitri S. Pavlichin, HoJoon Lee). Corresponding author Hanlee P. Ji, Division of Oncology, Department of Medicine – Stanford University School of Medicine, CCSR 1115, 269 Campus Drive, Stanford, CA 94305-5151.

## Abstract

K-mers are short DNA sequences that are used for genome sequence analysis. Applications that use k-mers include genome assembly and alignment. Despite these current applications, the wider bioinformatic use of k-mers in has challenges related to the massive scale of genomic sequence data. A single human genome assembly has billions of these short sequences. The sheer amount of computation for effective use of k-mer information is enormous, particularly when involving multiple genome assemblies. To address these issues, we developed a new k-mer indexing data structure based on a hash table tuned for the lookup of k-mer keys. This web application, referred to as KmerKeys (https://kmerkeys.dgi-stanford.org/), provides performant, rapid query speeds for cloud computation on genome assemblies. We enable fuzzy as well as exact k-mer-based searches of assemblies. To enable robust and speedy performance, the website implements cache-friendly hash tables, memory mapping and massive parallel processing. Our method employs a scalable and efficient data structure that can be used to jointly index and search a large collection of human genome assembly information. One can include variant databases and their associated metadata such as the gnomAD population variant catalog. This feature enables the incorporation of future genomic information into sequencing analysis.

## INTRODUCTION

The comprehensive genomic analysis enabled by whole genome sequencing **(WGS)** is generating a wealth of features. These include genetic variants of different classes such as structural variants **(SVs)**, diploid haplotypes of increasing size and complete sequence assemblies of individual chromosomes (1,2). Ultimately, these studies will substantially broaden our understanding of the full range of human genetic diversity and facilitate the discovery of genetic factors that influence disease susceptibility (3,4). These genomic features are described using different formats and distinct data types. Citing an example, catalogs of variants derived from population studies include the Genome Aggregation Database **(gnomAD)** (5), ClinVar (6) and The Cancer Genome Atlas **(TCGA)**. These variant resources provide their data across a variety of formats such as FASTA/Q, VCF and BED. To study these variants, it is critical to aggregate and organize this genomic data in a readily accessible fashion. Importantly, annotation of genomic features requires a format that can be related to different genome assemblies.

Catalogs of genetic variants require links to the human reference genome. With the completion of the International Human Genome Project, the current reference has frequently been updated (7-9). However, the current reference was constructed using the genome sequence of a small number of individuals and as a result, does not account for many genomic features across the breadth of human genetic diversity. Addressing this limitation, there are ongoing projects that involve constructing a pangenome reference derived from a broader sampling of the human population (10,11). These aggregated reference assemblies require the sequence analysis of hundreds of individuals (10,12). Linking human variant catalogs to this next generation of human reference genomes at this scale poses a significant challenge in terms of accessibility for genetic researchers.

K-mers are short lengths of sequence, typically in the tens of bases. K-mer analysis methods are appealing in their conceptual simplicity, because these short sequences can be readily manipulated and compared among different sequence data sets. K-mer-based tools have a variety of different functions that include: enumeration (13,14), read filtering (15), evolutionary distance estimation (16), metagenomics (17), and RNAseq analysis (18). The majority of applications are geared towards mapping sequences from FASTA/Q files. Beyond mapping, k-mers have specific advantages for organizing and querying sequence databases; one can index genomic data, facilitate the organization of these data sets and offer highly efficient querying of large collections of genomic sequence data.

Along these lines, we developed a website data resource (https://kmerkeys.dgi-stanford.org/) that indexes k-mer sequences for reference assemblies and links them to variant catalogs. Referred to as KmerKeys, this framework enables the user to include useful metadata such as genomic coordinates, counts, pointers to datasets and their inclusion of genetic variants. Moreover, our index incorporates commonly used genomic data formats (FASTA/Q, VCF, and BED). With these analytics, our website provides two special features: i) rapid, fuzzy searches from large assemblies such as the human reference genome and ii) representation of variants. First, one can conduct computationally efficient, fuzzy searches for a specific short sequence from a given reference. Our method retrieves neighboring short sequences that are a small edit distance away. Fast fuzzy searches enable approximate search in the space of k-mer-annotated variants and provides a uniqueness score that defines its representation from a given assembly. Second, a k-mer-based representation of genetic variants can easily be linked to a variety of metadata features.

With our approach, we indexed two human reference assemblies together with all common exon-based variants in gnomAD (version 2) (19). This first assembly was the established GRCh38 reference and the second was a telomere-to-telomere **(T2T)** haploid assembly of all 23 chromosomes in the CHM13 cell line, a hydatidiform mole which is an abnormal haploid outgrowth from a female egg (20). We used the T2T assembly as a haploid comparison data set. A query sequence is specified either as a text string or as genomic coordinates. Then, our web portal (https://kmerkeys.dgi-stanford.org/) returns the number and locations of all exact or approximate k-mer matches in GRCh38/T2T and associated variants in gnomAD. Approximate matches differ from the query by at most a user-specified maximum edit distance; gnomAD queries can be restricted to only sufficiently frequent variants, only indels, or using other filters. The web portal performs thousands to millions of such queries per second, depending on parameter settings and rapidly provides results.

## MATERIALS AND METHODS

### KmerKeys web application

KmerKeys is our web portal (https://kmerkeys.dgi-stanford.org/) deployed as a public cloud-hosted service on Amazon Web Services **(AWS)**. We generated a cloud-based index of two genome assemblies (GRCh38 and a T2T assembly) and the exonic variants in gnomAD (version 2) and made these indices publicly available in a web application **(Figure 1)**. All indices on KmerKeys are searchable either by sequence in FASTA-style input or by a set of intervals within a selected dataset in genomic coordinates. Both exact and fuzzy queries are supported, the latter performed within some maximum allowed number of mismatches. When querying by sequence, an input string is decomposed into its constituent k-mers in a sliding window fashion, which are then individually queried against the hash table. When querying by coordinate, we first retrieve the sequence at the specified coordinates, and then query its constituent short k-mers.

**Figure 1.**
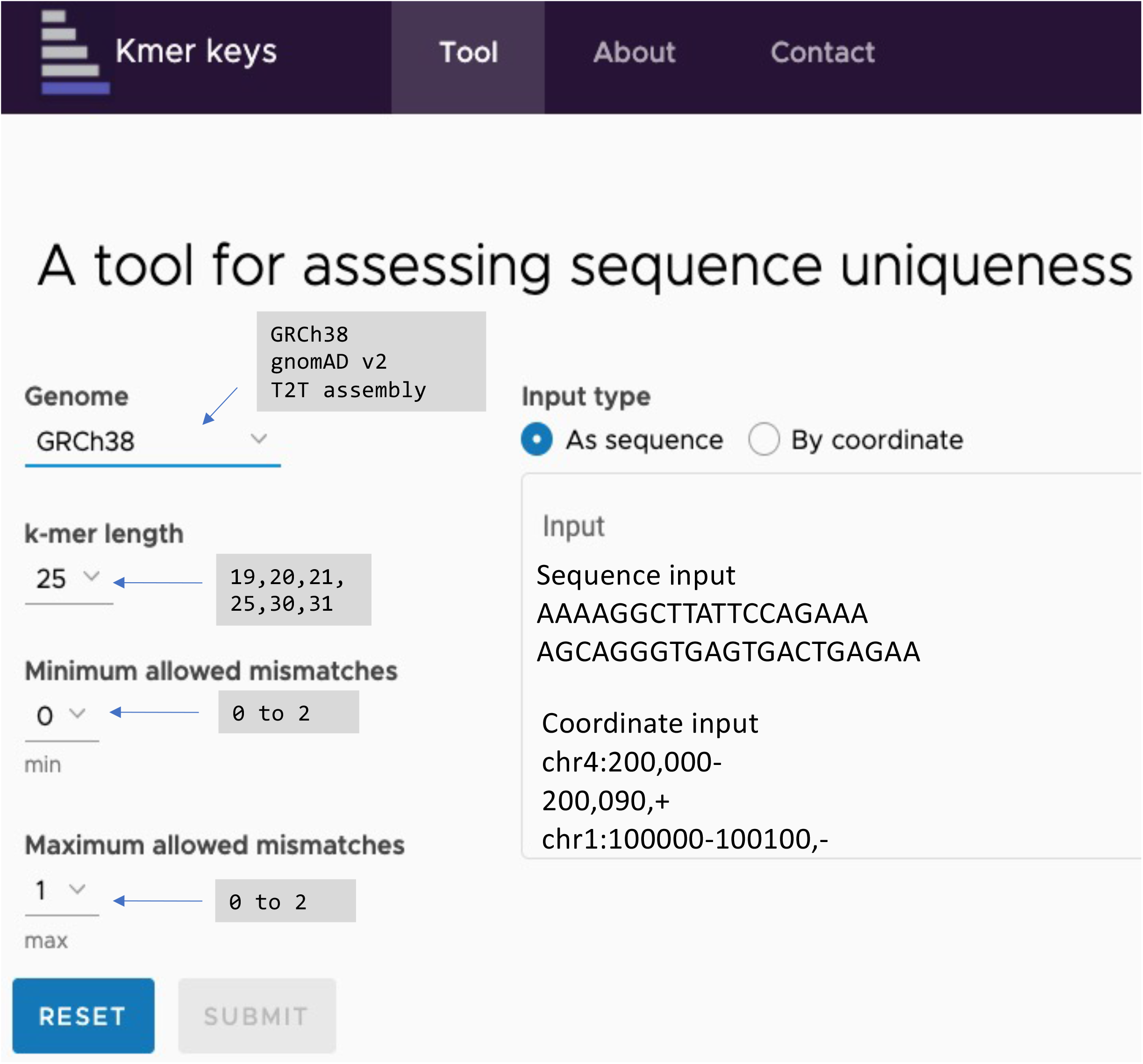
Web application of KmerKeys. There are three indices available to query; i) GRCh38, ii) gnomAD v2, and iii) T2T assembly of CHM13. Users can query these indices based on k-mers from length 19 to 31, and using either sequences or coordinates as input. Results will be generated with the user’s choice of fuzzy search by setting minimum/maximum allowed mismatches.

### Data structure

The data structure involves a hash table associating k-mer keys with arbitrary values. This method is optimized for query speed. This data structure is related to quotient filters **(QF)** (21), the counting quotient filter (22), and similar data structures employed for storing short sequences (23,24). Similar to these other studies, our data structure has the advantage of high cache locality due to a linear hash collision resolution scheme and a compact representation of its key elements.

We use a fast, invertible hash function, amounting to a single multiplication both to generate and remove the hash. This approach is based on a simpler design than QF-like data structures to store and lookup keys. It eliminates the need for metadata bits and comparisons used in such data structures in favor of an approach relying on a few arithmetic operations. Our method introduces an overflow table populated by keys that are not inserted after a prescribed maximum number of hash collisions. An overflow table is an extra complication relative to other QF-like schemes. However, KmerKeys is designed to maintain a high speed performance by limiting the impact of overpopulated portions of the hash table and simplifying the k-mer count function (see **Supplementary Methods 1**). The use of an invertible hash function makes our data structure a lossless representation of the k-mer keys. Thus, one can recover any input from a given table.

Some k-mer counting tools such as Squeakr (24) use an invertible hash function applied to k-mers. In contrast, KmerKeys uses a single multiplication operation to both hash and unhash. In Squeakr, its inverse is more complicated since their hashing does not involve multiplication while their unhashing does. This requires more computational steps than the function we use in KmerKeys. Further, our data structure allows the keys to be stored in two possible orders: 1) sorted, or 2) approximately sorted. Concretely, for order item 2, approximate sorting means that two keys are separated by at least one empty bucket in the hash table in a sorted order. However, no such guarantee is made for two keys not separated by an empty bucket. In fact, order item 1 (sorted) simplifies some operations on multiple tables (like set union, intersection, and difference) relative to a random order; however, inserting a new key may require previously inserted keys to maintain the sorted order. Unlike order item 1, order item 2 (approximately sorted) guarantees that previously inserted keys never change address in the table, which is a useful property if a key’s address is used to access another table, since otherwise we would need to reorder the other table when key addresses change. Nonetheless, order item 2 still permits set operations to be computed almost as efficiently as order item 1, at the expense of a more complicated implementation. This property simplifies the construction of associative arrays such that our hash table stores their keys, while mostly maintaining the computational advantages of having a precise soring order (see **Supplementary Methods 2**).

### Hashing k-mers

We represent a k-mer by the last 2k bits of an unsigned integer (we support up to k = 64 using unsigned 128-bit integers). We let the nucleotide ‘A’ correspond to the bits ‘00’, ‘C’ to ‘01’, and so forth. Then we might have a 31-mer like:

A A C G A T C G A C G T G C C C T T A C G C A A A G C T T A T (31-mer) 00000110001101100001101110010101111100011001000000100111110011 (binary)

Quotient filter-like data structures treat the binary k-mer representation as a number x in the range [0, 4^k-1] and allow the quotient of x upon division by another integer N to be the “home address” of x in a table. This scheme results in many hash collisions when a subset of k-mers share a common prefix, as is commonly the case in genomic data. To reduce the frequency of shared prefixes, we apply a multiplicative hash function to the binary representation x of a k-mer:

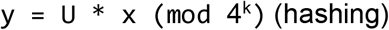

and inverse

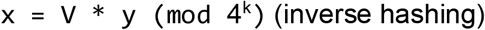

where U and V are integers such that U * V = 1 (mod 4^k). By using Bézout’s identity and the extended Euclidean algorithm, we can always find a number V such that V * U = 1 (mod B) whenever U and B have no prime factors in common; that is, multiplication by V inverts (unhashes) multiplication by U. This hash function is an automorphism on the set of k-mers viewed as the set, invertibly mapping “unhashed” to “hashed” k-mers.

When B = 4^k^, any odd number U works, but we choose U to be an odd integer near 4^k^ / ϕ, where ϕ = (1 + √5)/2 ≈ 1.62 is the golden ratio. This choice, known as Fibonacci hashing, increases the expected spacing of the hash values of consecutive keys. Both the hashing and unhashing operations correspond to a single integer multiplication and bitwise AND to compute the mod (since 4^k^ is a power of 2). For example, for k = 30 we choose U = 712544676207699905 and V = 201872644167657537, where 4^30^ / U ≈ ϕ and U * V = 1 (mod 4^30^).

### Constructing and querying the hash table

Our hash table has a number of important parameters that include the sequence of length K, the number of slots N and an integer H which is used to determine the maximum distance of a key from its “home slot”. Keys farther than H slots from their home slot are stored in an overflow table (see **Supplementary Methods 3**). Additionally, we add trait-like parameters allowing for different behaviors of our data structure -- whether keys are stored in a sorted order and whether duplicate keys are allowed.

A basic quotient filter idea is to associate each k-mer with a “home slot”: if x is a k-mer represented as a 2k-bit integer, then x can be uniquely represented as x = q * L + r, where q is the quotient of x upon division by L, and r is the remainder. Then q is the home address of x in our hash table, where L is the number of k-mers with the same home address (namely, L = ⌈ 4^k^ / (N – H)⌉, where ⌈ · ⌉ denotes the ceiling function. Typically, N is much larger than H, so L ≈ 4^k^ / N.

The content of the home slot is an invertible function of the remainder r (requiring fewer than all 2k bits to represent x), so that a k-mer x can be reconstructed from its address q and the contents stored at q. The particular invertible function we use is L - r (this has the advantage that empty slots store the value 0, since r is an integer between 0 and L – 1). In the case of a hash collision such that two k-mers have the same home slot q, we increment the slot address q until we find the nearest empty slot h slots away. This is linear hash collision resolution. If h > H, then we store the key in the overflow table. Otherwise, we store L * (h + 1) - 1 in the slot.

### Associating metadata with a k-mer

The k-mer storage address is used as an index for one or more separate metadata tables. In our web application for two assemblies, these metadata tables store the number of appearances and the assembly coordinates of each k-mer. Due to large entries (more than billions), some multiple hashings occupy the same spot, which we term a “hash collision”. In other word, this happens when linear hash collision resolution fails to find an empty slot after a maximum number of attempts. To prevent this, the k-mer metadata is written to an external metadata overflow table when an overflow occurs by attempting to insert a k-mer into our hash table. This overflow table is implemented using any convenient dictionary-like data structure (like a dict in a Python implementation) and is assumed to be accessed rarely. Otherwise, we rehash the table contents to a larger hash table. So long as the overflow table is accessed rarely, this choice has a small effect on the overall performance of our data structure. This metadata structure is also used in annotating variants for comparisons among different genome assemblies.

### Web implementation

The front-end is a web app created using Angular (https://angularjs.org/). The front-end interfaces with a computational back-end running on a separate server, an AWS EC2 instance with sufficiently large memory to support k-mer indices for GRCh38, a T2T genome, and gnomAD v2 for multiple values of K. Queries submitted via the front-end are sent to the back-end, which generates a response returned via the front-end. Moreover, query responses can be downloaded via a link we provide to the user, valid for one hour after query completion. Occasionally, query responses are too large to be conveniently printed on the front-end. In this case, we print an abbreviated response to the front-end; users can use the download link to download the full, unabbreviated response.

## RESULTS

### Overview of KmerKeys

KmerKeys is a performant data structure that associates arbitrary metadata with k-mer keys, allowing for large query speed and fuzzy search. In the hash table of KmerKeys, the bipartite variant graph has billions of k-mers (circles) and millions of locations (squares) **(Figure 2A)**. For GRCh38 and the T2T assembly, KmerKeys has the hash table of all k-mers from both genomes as keys and associates them with the following metadata: i) the frequencies of the k-mer in GRCh38/T2T assembly and ii) the k-mer location(s) in GRCh38/T2T assembly. Therefore, each location will be associated with a given short sequence at that position, but also could be linked to multiple locations if the k-mer appears multiple times. KmerKeys allows the user to search these indices using these short sequences and the coordinates of assemblies.

**Figure 2.**
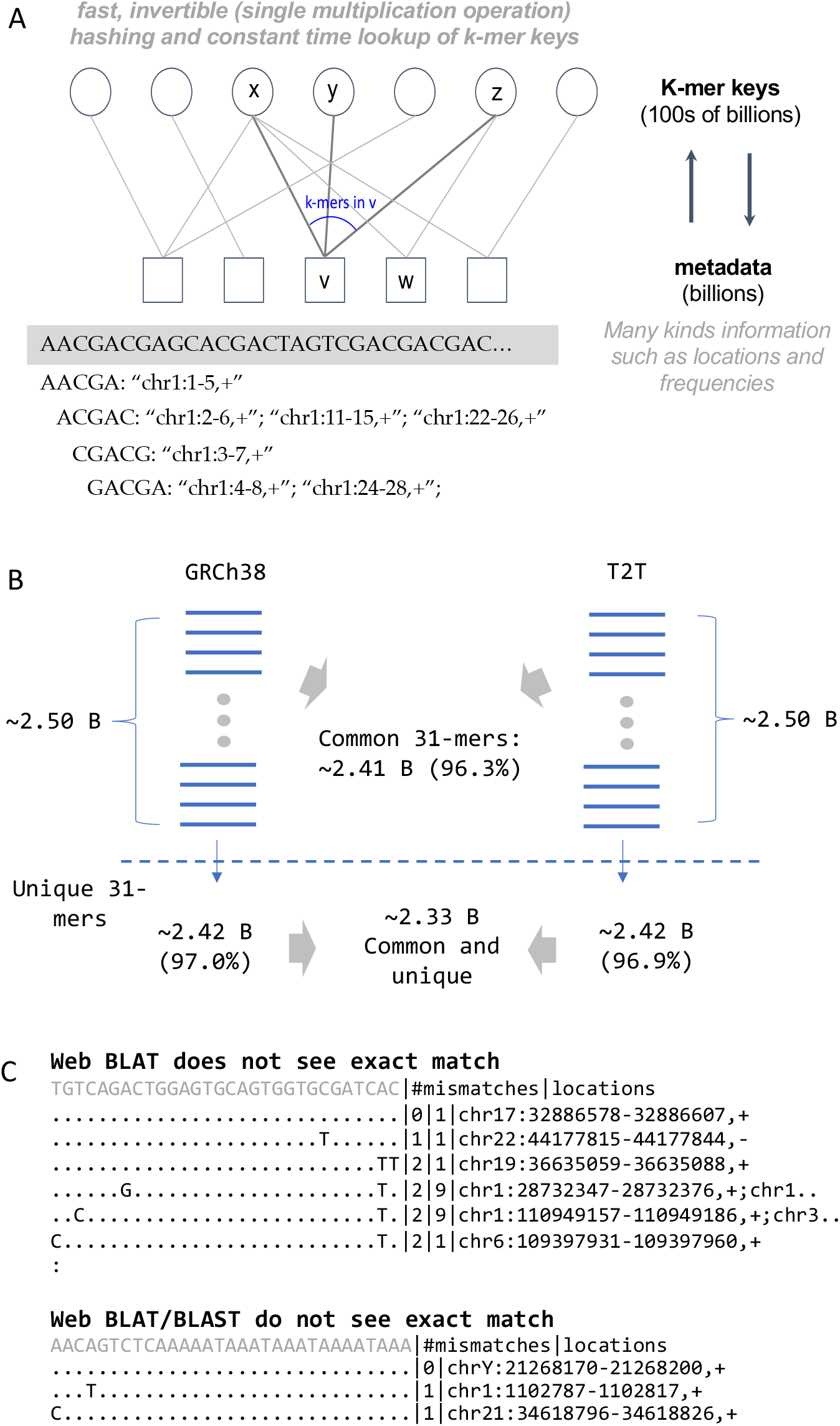
Overview of KmerKeys. A) Data structure of KmerKeys. The bipartite variant graph structure associates each k-mer key (circle) with its metadata (square). B) Index of 31-mers for GRCh38 and T2T assembly. Each of these genomes contains approximately 2.5 billion 31-mers. Most 31-mers occur only once within each genome (i.e. unique) and most 31-mers are common between the two genomes. C) Example k-mers with unique exact matches identified by KmerKeys but not found by the web versions of BLAT/BLAST.

We developed KmerKeys in the Julia programming language (25,26). The primary benefit of Julia is its level of language expressiveness and concision similar to Python, enabling rapid prototyping and experimentation without sacrificing much performance relative to compiled languages like C and C++. Thus, using Julia enabled us to prototype and release a performant version of our tool in the same language, which accelerated development.

The KmerKeys website was designed and optimized for high query speed, anticipating that indices would be rarely constructed and frequently queried. This choice of tradeoff is suited for a cloud-based shared resource where the same memory and compute resources are shared by multiple users, and the architecture can straightforwardly scale to ever more assemblies and larger datasets. All visitors’ queries to the KmerKeys public resource use the same pool of memory and threads, thus reducing the average cost per query. The web application also allows anyone without a computational background to readily access our tool.

### Operations of KmerKeys

Currently, KmerKeys hosts indices of two whole genome assemblies – GRCh38 and the T2T assembly of CHM13. We used six different lengths of k-mers ranging from 19 through 31 – specifically, 19, 20, 21, 25, 30 and 31. For example, KmerKeys indexed ∼2.5 billion distinct 31-mers from both GRCh38 and the T2T assembly, and ∼2.41 billion 31-mers (96.3%) were present in both GRCh38 and the T2T assembly **(Figure 2B)**. Most 31-mers (97%) are unique, appearing once in GRCh38 or the T2T assembly and thus associated with a unique location. The same trends were observed for values of k between 19 and 31. The k-mers that are present multiple times in GRCh38 indicate sequences that occur in multiple locations. Among the common 31-mers between the two assemblies, 97% of them are unique, appearing once in GRCh38 and the T2T assembly respectively. These unique 31-mers present in both assemblies have potential use as Sequence Tagged Sites **(STS)** which have a single occurrence in the genome and a known location and base sequence (27).

To increase the performance, we employed a number of mathematical concepts previously unexploited in the k-mer indexing setting. They include an invertible Fibonacci hash function together with linear hash collision resolution and a quotient filter-inspired bitpacking scheme. Together, these features offer fast (constant expected time) queries that leverage memory caching for speed, bitpacking to reduce space and a simpler implementation than related data structures like the quotient filter. Further, we used this hash table to associate k-mers with metadata, thereby supporting optional memory mapping of values (the metadata) or the keys to reduce memory usage, and optimizing further for the setting of indexing locations and counts in a FASTA file. As a result, our implementation of this hash table supports millions of table lookups per second on a single thread.

KmerKeys offers accurate search capability that existing tools such as BLAST (28) and BLAT (29) do not. To note an example, KmerKeys identified k-mers appearing uniquely in GRCh38 which were not identified by BLAST or BLAT **(Figure 2C)**. We found that BLAT sometimes fails to find k-mers that exist at a unique location in GRCh38. For instance, TGTCAGACTGGAGTGCAGTGGTGCGATCAC (30-mer) contains no obvious low complexity sequences. However, BLAT reported several approximate matches, but not the unique exact match at chr17:32886578-32886607. In addition, the web versions of BLAT and BLAST failed to identify the unique exact match at chrY:21268170-21268200, AACAGTCTCAAAAATAAATAAATAAAATAAA (31-mer).

We randomly sampled 100,000 unique 20, 21, 30 and 31-mers in GRCh38 (that is, each k-mer occurs at exactly one position in GRCh38). Using BLAT, we failed to identify 1,318 of the 20-mers (1.32%), 1,350 of the 21-mers (1.35%), 435 of the 30-mers (0.435%) and 440 of the 31-mers (0.44%), respectively **(Supplementary File 1)**. We also found that the web-based BLAST (30), though not the standalone software, sometimes misses unique k-mers. The same 100,000 randomly sampled unique 31-mers were identified by standalone BLAST, but web-based BLAST fails to find 316 of the uniquely occurring 31-mers among 462 that were not found by BLAT.

These missed unique k-mers contain the over-represented (appears more than 1024 times) 11-mers and the vast majority of them would be masked as repeat elements by RepeatMasker. Interestingly, several of the missing k-mers were located in coding regions. For example:

- ATGTTTTATTTTCATGGGGC (20-mer), chr2:181522191-181522210, *ITGA4*
- CTCGCCTCCTGGGTTCAAGCAATTCTCCCG (30-mer), chr11:31681824-31681853, *EPL4*

To save computation time, BLAT and BLAST utilize 11-mers for the initial search for potential genomic regions where the actual sequence could be found. KmerKeys, on the other hand, simply indexes these short sequences of all k-mers as they are, thus it guarantees the completeness of its search results. KmerKeys also identifies all approximate matches along with their locations within a given number of mismatches. These features are not standard outputs to either BLAT or BLAST.

### Uniqueness of k-mers in the human reference genome (GRCh38) and the T2T assembly

Fuzzy, approximate searching provides a comprehensive landscape of uniqueness of k-mers in the human reference genome by length of K. Some of these may be a unique sequence and occur at a single coordinate for a given genome assembly. However, some sequences remain unique within one or two mismatches, which would make these k-mers useful for indexing single nucleotide variants and polymorphisms **(SNPs)**. The extent of uniqueness varies. In fact, most distinct 20-mers (>90%) are unique, but about 63% of them are not unique within 1 mismatch and about 99% are no longer unique within 2 mismatches. For 31-mers, 88.9% and 82.8% of 31-mers are unique within 1 and 2 mismatches, respectively **(Figure 3A)**. The information about neighbor k-mers is not easily retrieved by widely used tools such as Jellyfish (13) and KMC (14). However, our exact and fuzzy searches collect this type of information simply by retrieving the metadata that includes assembly coordinates and frequency. Importantly, this metadata includes information about neighbor k-mers.

**Figure 3.**
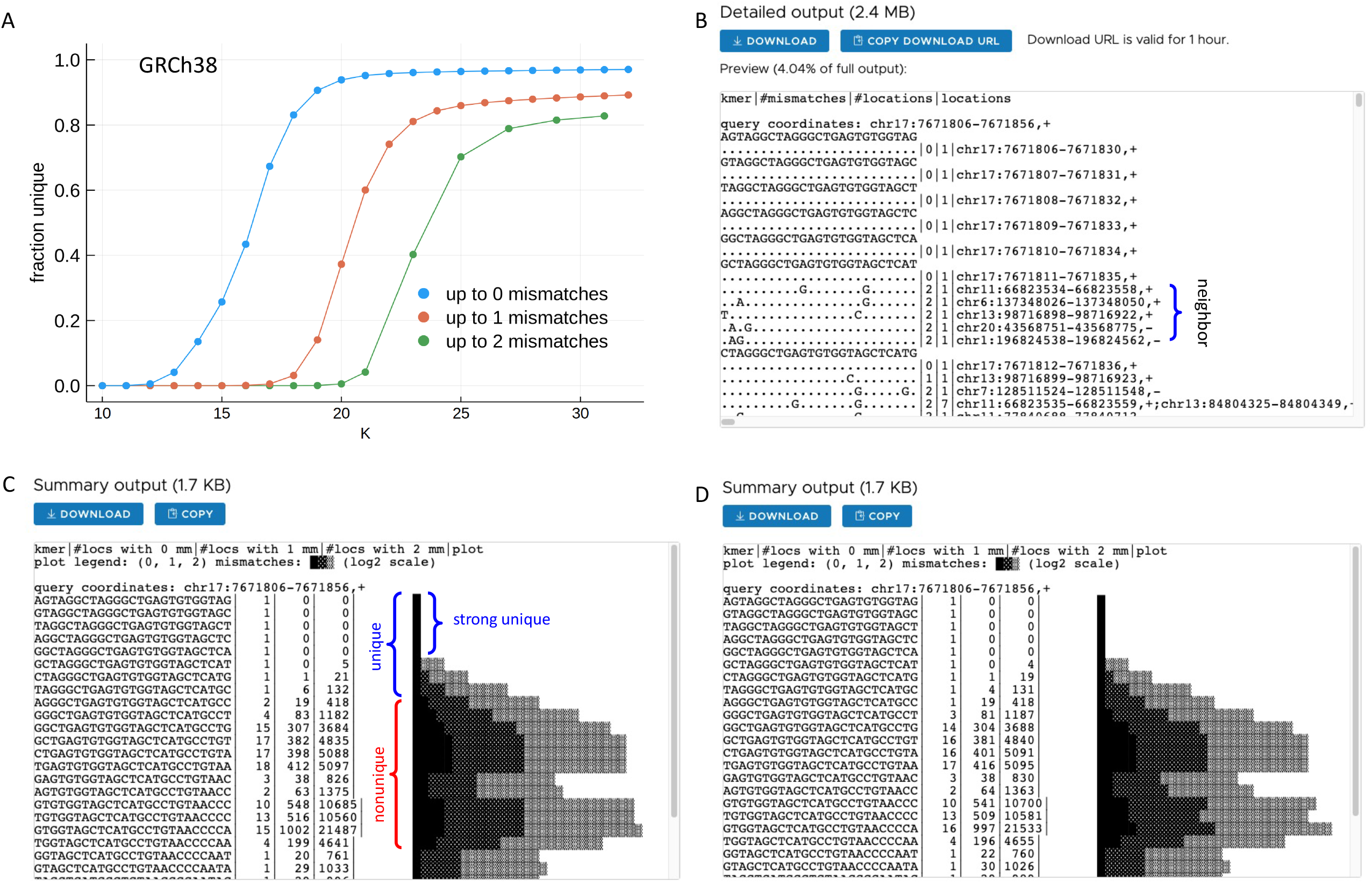
The uniqueness of k-mers measured by fuzzy search. A) Overall landscape of k-mer uniqueness within 2 mismatches by length of k-mers. Longer k-mers are more likely to remain unique up to 2 mismatches. B) Example of detailed output from a KmerKeys web application query of 25-mers in GRCh38 in an intronic region of *TP53*. The detailed output shows unique exact match k-mers as well as neighbor k-mers, with their coordinates, frequencies, and any mismatched nucleotide positions. C) Example of summary output from a KmerKeys web application query of 25-mers in GRCh38 in an intronic region of *TP53*. The summary output displays each k-mer sequence in sequential order with counts of k-mers at each edit distance up to 2 mismatches, along with a plot that visually displays those counts. D) Example of summary output for the same query in the T2T assembly.

Knowing the uniqueness of any sequence is critical information for a range of applications. This property is characterized by the following: i) the frequency of a given sequence at a specific assembly coordinate, ii) the number of neighbor k-mers that are a small edit distance away, iii) the frequency of neighbor k-mers with their locations and iv) the positions of the mismatching nucleotides on the neighbor k-mers. **Figure 3B** shows an example of detailed outputs from the query of the coordinates, chr17:7671806-7671856 of GRCh38, which is located within the intron between exons 5 and 6 of *TP53*. Identical nucleotides are indicated by a dot (.) and different nucleotides are shown at their positions. The example search result showed that the k-mer sequences from the first 5 positions are unique within 2 mismatches while the sequence at the 6^th^ position has 5 neighbor sequences with 2 mismatches. For instance, the 25-mer at the 6^th^ position is identical to the 25-mer at chr6:137348026-137348050 except for two mismatches: i) A instead of T at the 3^rd^ base position and ii) G instead of A at the 19^th^ base position. The 25-mer at the 7^th^ position has one neighbor sequence with 1 mismatch and 12 neighbor sequences with 2 mismatches (not all of them displayed). The k-mer at the last visible row could be matched to the query 25-mer with 2 substitutions (A to G at the 10^th^ base position and A to G at the 18^th^ base position). This 25-mer appears at 7 locations in the genome, including chr11:66823535-66823559 and chr13:84804325-84804349.

In addition to the above detailed output, KmerKey provides a summary output which allows the user to rapidly review the landscape of sequence uniqueness across a region of interest. The summary output shows the overall frequency of all query k-mers grouped by edit distance in the following format: i) the sequence of the k-mer, ii) the number of locations with an exact match, iii) the number of locations with an edit distance of 1 and iv) the number of locations with an edit distance of 2. Further, we provide a visual summary plot directly adjacent to the table. As seen in **Figure 3C**, the upstream region of interest contains unique sequences while the downstream region of interest contains non-unique sequences in GRCh38. Similar trends were observed in the T2T assembly although there are minor differences **(Figure 3D)**. As shown in **Figure 3C and D**, the 9^th^ 25-mer is not unique in GRCh38 but is unique in the T2T assembly.

KmerKeys has two outputs: i) a summary output and ii) a detailed output. To enable efficient online querying, the detailed output displays only the first 1000 lines. To allow the user to obtain results for queries that extend beyond 1000 lines, files are written to an AWS S3 bucket with a download link generated for users which is available for one hour.

### K-mer-based representation of variants

We developed a method to represent genetic variation that includes single nucleotide variants **(SNVs)** and insertion deletions **(indels)**. A variant is represented as a collection of k-mers that are unique to the genome of interest relative to the reference genome lacking the variant. For a single base pair substitution, this is the set of k-mers overlapping the substituted base pair. For short indels, this is the set of k-mers spanning the insertion or deletion **(Figure 4A)**. Importantly, this representation of a variant does not depend on a reference coordinate system. That being said, we can associate assembly coordinate metadata or any other kind of metadata, like clinical information. A variant is depicted as a square node in **Figure 2A**, thus incurring no loss in descriptive power relative to using the VCF format to describe variants.

**Figure 4.**
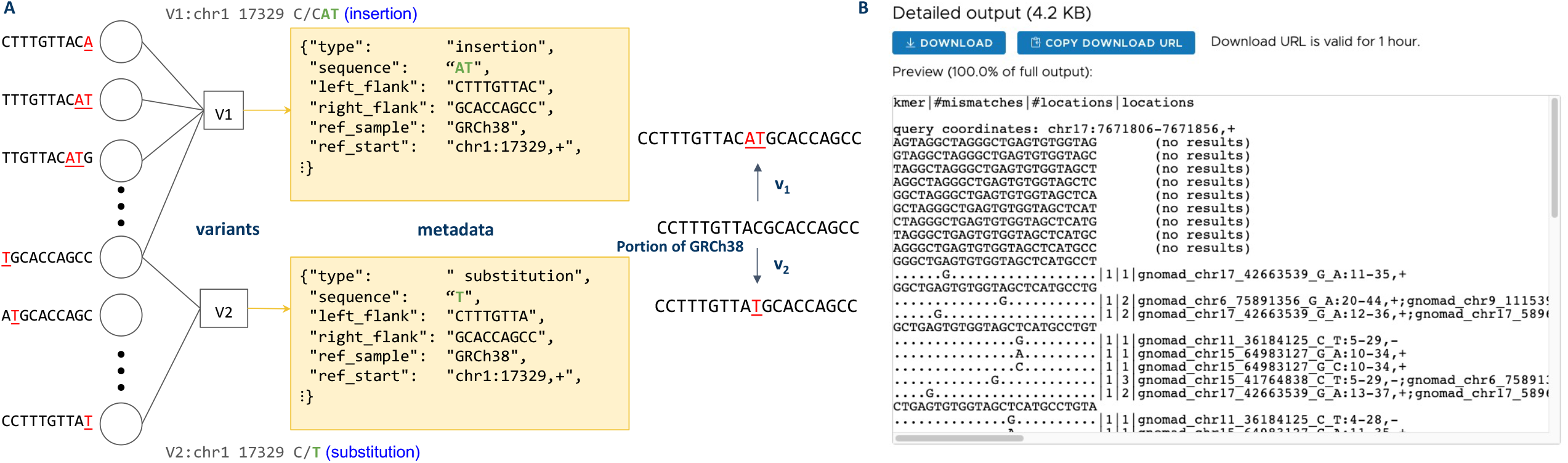
Association with metadata of variants. A) k-mer representation of variants. In the bipartite variant graph, each k-mer key (circle) is a k-mer generated by a variant in the GRCh38 sequence. The k-mer keys are associated with metadata (square), which includes the variant coordinate, sequence, type, and other useful information. B) Example of detailed output from a KmerKeys web application query of 25-mers in gnomAD v2 in an exonic region of *TP53*.

As an example, consider a particular variant: chr1:17329 C/CAT -- an insertion of the novel sequence AT following the ‘C’ at position chr1:17329 in GRCh38. Then the reference sequence in GRCh38, lacking the variant, is:

> …CCTTTGTTACGCACCAGCC…

And a “patched” version instead containing the alternate allele is:

> … CCTTTGTTAC<u>AT</u>GCACCAGCC…

where the inserted sequence is underlined. This set of novel k-mers overlaps any part of the variant allele, as shown in **Figure 4A**. Variant metadata can also include the variant coordinates in the reference genome as well as adjacent k-mers that are not affected by the variant, labeled “left_flank” and “right_flank” in **Figure 4A**. These extra metadata features make it straightforward to reconstruct a VCF-style representation of a variant from its graphical representation, and to map the variant to new assemblies.

By associating k-mers with variants, we enable fuzzy sequence-based search of a collection of variants. This function has some novel properties. For example, with a VCF file one can query a set of variants based on genome coordinates. However, it is less easy to do so based on the actual sequence containing the variation. Using a conventional approach, sequence-based variant querying requires one to first align the sequence to a reference genome and then use some other method to describe the set of variations. In contrast, our method permits associating short DNA sequences directly with k-mers and their metadata. A fuzzy search is achieved by querying all sequences within a given Hamming radius of a query sequence.

Fuzzy searching of k-mer-represented variants also enables robustness to the case of ambiguous variant definitions. For example, a substitution “AC/GG” may be represented as a pair of one-base-pair substitutions or a single two-base-pair substitution. By returning all approximately matching variants within some tolerance one can identify a maximum number of mismatches per k-mer.

### Population variant searching from gnomAD

We demonstrated the extensibility of our data structure for population-based genetic variation. This involved generating an index of all exonic variants in gnomAD v2 through our k-mer-based variant representation. Basically, the set of k-mers for a given length k overlapping the substituted base pair or spanning the insertion or deletion were associated with the coordinates based on GRCh38 **(Figure 4A)**. A collection of variants represented in this way corresponds to a bipartite graph, with k-mers on one side and variants on the other denoted as circles and squares in **Figure 2A**. Some k-mers may be associated with multiple variants, with fewer k-mers associated with multiple variants as the k value increases. In general, a k length of 19 to 31 is likely to be associated with only a single SNV across an entire human genome as we show in **Figure 3A**.

KmerKeys linked 17,119,203 variants in gnomAD with 523,498,431 31-mers. Out of 523,498,431 31-mers, 518,308,269 (99.0%) are unique. In contrast, 513,901,185 (98.2%) were not found in GRCh38 which indicates that they can be mapped unequivocally to a particular variant. In fact, 16,923,518 (98.9%) variants from gnomAD v2 have at least one unique 31-mer, which were not found in GRCh38.

The bipartite variant graph has hundreds of millions of k-mers (circles) and millions of variants (squares) for the exonic variants of gnomAD (v2) available to query on our web portal. Users can examine if their region of interest contains the variants reported in gnomAD v2 by querying a sequence or coordinates based on GRCh38. In **Figure 4B** we demonstrate an example using the gnomAD variants found in the last exon of *TP53*, chr17:7669515-7669660. All the 25-mers within this region except 10 bp of the upstream portion overlap with at least one variant. This information can also be obtained by providing the sequence from chr17:7669585-7669671. It is important to note that none of the k-mers associated with variants are present in GRCh38. The fuzzy search function makes it possible to demonstrate how variants with unique k-mers from other genomes can be mapped back to the reference. This feature is unavailable in any other public tool of which we are aware. This function could provide useful information about whether 20-mers of interest could be unique in other individuals.

### Comparison of k-mers and the arrangements from different assemblies

We demonstrated how the KmerKeys approach could be used for making comparisons between different genomes. This type of analysis facilitates the identification of features such as structural variants. For our current version, we provide two assemblies indexed by KmerKeys; GRCh38 and the T2T assembly of CHM13. KmerKeys enables visual inspection of possible genomic rearrangements by examining the arrangement of k-mers based on their coordinates in each assembly. Here, we identified four structural variants in CHM13 of the T2T assembly reported from Audano *et al*. (15): i) an inversion near chr19:38765000-38795000 on GRCh38, ii) a deletion near chr22:17275000-17300000 on GRCh38, iii) an insertion near chr18:67650000-67675000 on T2T and iv) a duplication near chr16:30190000-30290000 on GRCh38.

For each unique k-mer in GRCh38, we determined its position and strand alignment in the T2T assembly. Subsequently, our results are visualized using matrices comparing overlapping segments of the two assemblies. When an inversion occurs, the order of k-mers is reversed relative to the reference genome **(Figure 5A)**. When a deletion occurs relative to the reference genome, the pairs of k-mers that are distantly positioned in the reference genome are brought close together **(Figure 5B)**. For the case of an insertion, pairs of k-mer that are next to each other in the reference genome become separated by a large distance **(Figure 5C)**. The set of k-mers – unique in GRCh38 - involved in a duplication appear multiple times in T2T **(Figure 5D)**. Plotting the relative location of each k-mer and its strandedness clearly reveals the pattern of structural variants.

**Figure 5.**
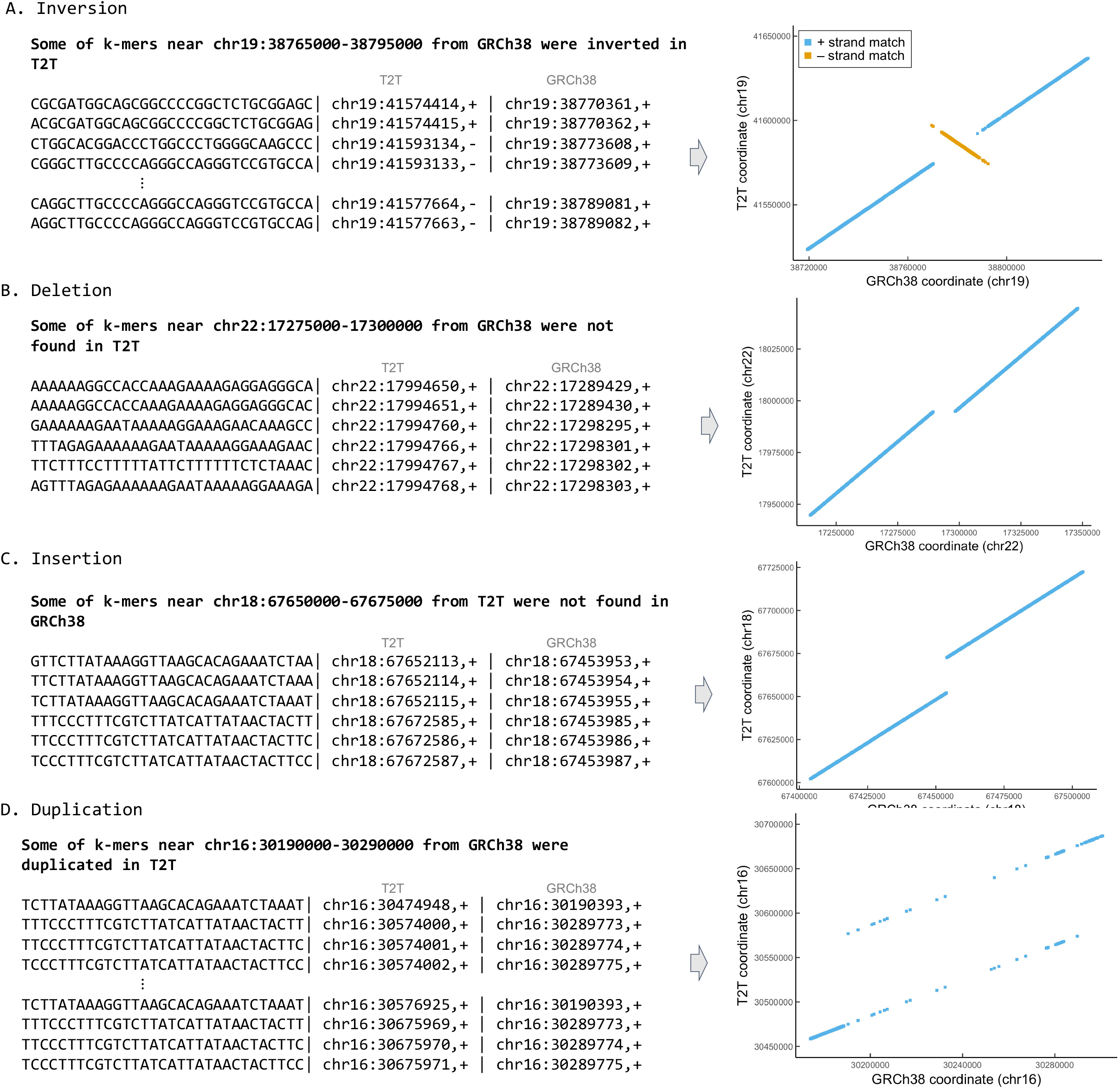
Visualization of four structural variants in T2T with respect to GRCh38. The position of each k-mer in each genome assembly was plotted in a dot matrix to reveal the pattern of an inversion (A), deletion (B), insertion (C), and duplication (D). Perfect matches are indicated by a straight diagonal line that is plotted from the two assemblies. Offsets and interruptions of the diagonal are a result of structural variation in which unique k-mers within an assembly are shifted in their positions. This feature allows for rapid visual recognition of rearrangements.

We demonstrated that the indexing of two assemblies enables visual inspection of potential structural variations by plotting the set of k-mers using coordinates from two different assemblies. For structural variations, we can generalize this approach by defining variants as functions of multiple k-mers. This representation also avoids having to precisely define start and end positions of structural variants, which frequently make it difficult to compare structural variants from multiple samples. Future studies will establish a k-mer-based grammar for defining other structural variations.

## DISCUSSION

In this study, we describe KmerKeys, a web data application that provides k-mer-based querying of human genome assemblies. For this application, we achieved the following: 1) we developed a data structure that efficiently and accurately associates arbitrary metadata with k-mers, 2) we devised a k-mer-based representation of variants that allows lists of variants to be jointly indexed with assemblies and primary sequencing, and 3) we launched a web application demonstrating the above, allowing users to query the locations and counts of k-mers in two whole human genome assemblies and exonic gnomAD v2 variants. Through these functions, we extended the ability to integrate, index and search large collections of diverse genomic types and formats (FASTA/Q, BED, VCF) beyond what is currently easily possible with existing tools, focusing on large cloud deployments optimized for query speed. Additionally, the search speed of our data structure is fast enough to enable fuzzy searches. This feature allows for tolerance of varying numbers of mismatches within a k-mer key. One potential application is finding near-matches in the cases of sequencing primer design, CRISPR/Cas9 target design and known variants in the populations.

The KmerKeys search function highlights a conceptual difference between our variant representation and how variants are generally reported. Traditionally, variants are described in terms of a reference and an alternate allele. This alternate allele involves a substitution, deletion, insertion, etc. that alters the sequence. The exact genetic mechanism can vary for any given variant which may change the way the variant is classified. Our approach describes these variants only in terms of an observed sequence (specifically, its k-mer content). Thus, a variant becomes associated with a k-mer key and this feature enables rapid comparisons among multiple genomes.

Our approach has some distinct features compared to genome graph representations of assemblies. In the latter approach, a genome graph is constructed to represent multiple variants in a joint way, with individual variants not easily separable from a single global structure. Using KmerKeys, the variants are naturally separate entities that can be combined straightforwardly into new collections. Our approach can use filtering (e.g. those variants whose metadata is associated with a particular condition) or set operations on existing collections (e.g. variants in gnomAD v2 but not in ClinVar). These features avoid retaining information associated with a larger-than-necessary genome graph that contains many other variants that are not of interest to the user. KmerKeys can also be used to visualize structural variations. This process involves comparing the relative order of k-mers between different assemblies. To optimize the query speed meant a tradeoff in terms of greater memory requirements. However, we observed very high query speeds, especially for fuzzy search. Thus, our tool was well-suited for development as a shared web resource rather than requiring a local installation that requires local memory resources.

KmerKeys has the potential to be used for DNA primer design and CRISPR/Cas9 target design. Using its search function, one can identify primer candidates that have the potential for off-target sites. Further, our data structure could provide a framework for representing variants at the population level and in multiple genomes simultaneously. Our k-mer-based indices are designed with performance in mind. They are compact to minimize storage footprint and are designed to optimize query speed. We leverage cache-friendly hash tables, memory mapping, and optional parallelism to increase query performance. These features allow our approach to scale from personal machines to large computing clusters. For the task of k-mer counting alone, our approach is close to the state-of-the-art in terms of speed although we use more memory than other tools, while enabling significant novel indexing and querying functionality.

## Acknowledgements

We would like to thank Billy Lau for suggestions. We also thank Lucas Johnson for AWS setup and Jung Yoo for comments.

## Funding

This work was supported by National Institutes of Health grant [U01HG01096] and the Clayville Foundation.

## Conflict of Interest

Not applicable.

## Notes

https://kmerkeys.dgi-stanford.org/

### Competing Interest Statement

The authors have declared no competing interest.

## REFERENCES

1. Karczewski, K.J., Francioli, L.C., Tiao, G., Cummings, B.B., Alföldi, J., Wang, Q., Collins, R.L., Laricchia, K.M., Ganna, A., Birnbaum, D.P. et al. (2019) Variation across 141,456 human exomes and genomes reveals the spectrum of loss-of-function intolerance across human protein-coding genes. bioRxiv, 531210.

2. Stephens, Z.D., Lee, S.Y., Faghri, F., Campbell, R.H., Zhai, C., Efron, M.J., Iyer, R., Schatz, M.C., Sinha, S. and Robinson, G.E. (2015) Big Data: Astronomical or Genomical? PLoS Biol, 13, e1002195.

3. Van Hout, C.V., Tachmazidou, I., Backman, J.D., Hoffman, J.D., Liu, D., Pandey, A.K., Gonzaga-Jauregui, C., Khalid, S., Ye, B., Banerjee, N. et al. (2020) Exome sequencing and characterization of 49,960 individuals in the UK Biobank. Nature, 586, 749–756.

4. Dewey, F.E., Grove, M.E., Pan, C., Goldstein, B.A., Bernstein, J.A., Chaib, H., Merker, J.D., Goldfeder, R.L., Enns, G.M., David, S.P. et al. (2014) Clinical interpretation and implications of whole-genome sequencing. JAMA, 311, 1035–1045.

5. Karczewski, K.J., Francioli, L.C., Tiao, G., Cummings, B.B., Alfoldi, J., Wang, Q., Collins, R.L., Laricchia, K.M., Ganna, A., Birnbaum, D.P. et al. (2020) The mutational constraint spectrum quantified from variation in 141,456 humans. Nature, 581, 434–443.

6. Landrum, M.J., Lee, J.M., Riley, G.R., Jang, W., Rubinstein, W.S., Church, D.M. and Maglott, D.R. (2014) ClinVar: public archive of relationships among sequence variation and human phenotype. Nucleic Acids Res, 42, D980–985.

7. Schneider, V.A., Graves-Lindsay, T., Howe, K., Bouk, N., Chen, H.C., Kitts, P.A., Murphy, T.D., Pruitt, K.D., Thibaud-Nissen, F., Albracht, D. et al. (2017) Evaluation of GRCh38 and de novo haploid genome assemblies demonstrates the enduring quality of the reference assembly. Genome Res, 27, 849–864.

8. Lander, E.S., Linton, L.M., Birren, B., Nusbaum, C., Zody, M.C., Baldwin, J., Devon, K., Dewar, K., Doyle, M., FitzHugh, W. et al. (2001) Initial sequencing and analysis of the human genome. Nature, 409, 860–921.

9. International Human Genome Sequencing, C. (2004) Finishing the euchromatic sequence of the human genome. Nature, 431, 931–945.

10. Sherman, R.M. and Salzberg, S.L. (2020) Pan-genomics in the human genome era. Nat Rev Genet, 21, 243–254.

11. Li, R., Li, Y., Zheng, H., Luo, R., Zhu, H., Li, Q., Qian, W., Ren, Y., Tian, G., Li, J. et al. (2010) Building the sequence map of the human pan-genome. Nat Biotechnol, 28, 57–63.

12. Sherman, R.M., Forman, J., Antonescu, V., Puiu, D., Daya, M., Rafaels, N., Boorgula, M.P., Chavan, S., Vergara, C., Ortega, V.E. et al. (2019) Assembly of a pan-genome from deep sequencing of 910 humans of African descent. Nat Genet, 51, 30–35.

13. Marcais, G. and Kingsford, C. (2011) A fast, lock-free approach for efficient parallel counting of occurrences of k-mers. Bioinformatics, 27, 764–770.

14. Kokot, M., Dlugosz, M. and Deorowicz, S. (2017) KMC 3: counting and manipulating k-mer statistics. Bioinformatics, 33, 2759–2761.

15. Chen, S., Huang, T., Wen, T., Li, H., Xu, M. and Gu, J. (2018) MutScan: fast detection and visualization of target mutations by scanning FASTQ data. BMC Bioinformatics, 19, 16.

16. Deorowicz, S., Gudys, A., Dlugosz, M., Kokot, M. and Danek, A. (2019) Kmer-db: instant evolutionary distance estimation. Bioinformatics, 35, 133–136.

17. Wood, D.E., Lu, J. and Langmead, B. (2019) Improved metagenomic analysis with Kraken 2. Genome Biol, 20, 257.

18. Bray, N.L., Pimentel, H., Melsted, P. and Pachter, L. (2016) Near-optimal probabilistic RNA-seq quantification. Nat Biotechnol, 34, 525–527.

19. Francioli, L., Tiao, G., Karczewski, K., Solomonson, M. and Watts, N. (2018), gnomAD v2.1, pp. https://macarthurlab.org/2018/2010/2017/gnomad-v2012-2011/.

20. Audano, P.A., Sulovari, A., Graves-Lindsay, T.A., Cantsilieris, S., Sorensen, M., Welch, A.E., Dougherty, M.L., Nelson, B.J., Shah, A., Dutcher, S.K. et al. (2019) Characterizing the Major Structural Variant Alleles of the Human Genome. Cell, 176, 663–675 e619.

21. Bender, M.A., Farach-Colton, M., Johnson, R., Kraner, R., Kuszmaul, B.C., Medjedovic, D., Montes, P., Shetty, P., Spillane, R.P. and Zadok, E. (2012) Don’t Thrash: How to Cache Your Hash on Flash. Proc Vldb Endow, 5, 1627–1637.

22. Pandey, P., Bender, M.A., Johnson, R. and Patro, R. (2017) A General-Purpose Counting Filter: Making Every Bit Count. Sigmod’17: Proceedings of the 2017 Acm International Conference on Management of Data, 775–787.

23. Shokrof, M., Brown, C.T. and Mansour, T.A. (2020) MQF and buffered MQF: Quotient filters for efficient storage of k-mers with their counts and metadata. bioRxiv, 2020.2008.2023.263061.

24. Pandey, P., Bender, M.A., Johnson, R., Patro, R. and Berger, B. (2018) Squeakr: an exact and approximate k-mer counting system. Bioinformatics, 34, 568–575.

25. Bezanson, J., Edelman, A., Karpinski, S. and Shah, V.B. (2017). 1 ed, SIAM Review, Vol. 59, pp. 65–98.

26. Perkel, J.M. (2019) Julia: come for the syntax, stay for the speed. Nature, 572, 141–142.

27. Olson, M., Hood, L., Cantor, C. and Botstein, D. (1989) A common language for physical mapping of the human genome. Science, 245, 1434–1435.

28. Altschul, S.F., Gish, W., Miller, W., Myers, E.W. and Lipman, D.J. (1990) Basic local alignment search tool. J Mol Biol, 215, 403–410.

29. Kent, W.J. (2002) BLAT--the BLAST-like alignment tool. Genome Res, 12, 656–664.

30. Wheeler, D. and Bhagwat, M. (2007) BLAST QuickStart: example-driven web-based BLAST tutorial. Methods Mol Biol, 395, 149–176.

