## Supplementary Methods for "KmerKeys: a web resource for searching indexed genome assemblies and variants"

[†] Co-first authors

* Corresponding author

Hanlee P. Ji

Division of Oncology, Department of Medicine – Stanford University School of Medicine

CCSR 1115, 269 Campus Drive

Stanford, CA 94305-5151

### SUPPLEMENTARY METHODS

#### 1. Hash table

Our hash table T is parametrized by integer k (k-mer length), integer A (alphabet size; A=4 for the 4 nucleotides {A,C,G,T}), integer N (the number of slots in an array), and integer H, the maximum distance of a key from its "home slot" (defined below). Keys farther than H slots from their home slot are stored in an overflow table $T_{ov}$. Additionally, we add trait-like parameters allowing for different behaviors of our data structure: Boolean IsSorted (controls whether keys are stored in a sorted order) and Boolean AllowDuplicates (controls whether duplicate keys are allowed).

- Integer k: k-mer length
- Alphabet size A (A = 4 for the nucleotides {A,C,G,T})
- Integer H (≥ 0): the maximum distance of a key from its "home address" (defined below)
- Integer N (≥ H): the number of slots in an array of slots

Keys farther than H slots from their home slot are stored in an overflow table $T_{ov}$. Additionally, we add trait-like parameters allowing for different behaviors of our data structure:

- Boolean IsSorted – controls whether keys are stored in a sorted order
- Boolean AllowDuplicates – controls whether duplicate keys are allowed

Several parameters are derived from the parameter choices above, and are convenient to reference below:

- A pair of integers U and V such that U * V = 1 (mod $A^k$) and U is the nearest integer to $A^k$ / ϕ with no prime factors in common with $A^k$, where ϕ is the golden ratio: multiplication by U (resp. V) hashes (resp. unhashes) k-mer keys.
- Integer L = $\lceil A^k / (N – H) \rceil$, where $\lceil . \rceil$ denotes the ceiling function: L is the number of distinct k-mers with the same "home address" (defined below).

- Integer $B \geq \lceil \log_2(H * L + 1) \rceil$: the number of bits per slot in the hash table. In our implementation we round B up to a multiple of 8, so that a slot is some integer number of bytes.

The N slots of hash table T are addressed by address q in {0,1,…,N-1}. Let T[q] denote the numerical value stored at address q in T. Since B bits are used per slot, then T[q] is in $\{0,1,\ldots,2^B-1\}$.

Let x be a k-mer and let underlines denote hashed quantities, e.g. $\underline{x} = U * x \pmod{A^k}$ and $x = V * \underline{x} \pmod{A^k}$.

The slot at address q is either "empty" or corresponds to a hashed k-mer $\underline{x}_q$ (and unhashed k-mer $x_q$). Slot q is empty if T[q] = 0. If slot q is not empty (if T[q] ≠ 0) , then the corresponding stored (hashed) k-mer is computed as:

$$\underline{x}_q = q * L + (L - T[q])$$

The "home address" of (hashed) k-mer $\underline{x}$ is the unique non-negative integer q satisfying the equation:

$$\underline{x} = q * L + r$$

with r in {0,1,…,L-1}. That is, q is the quotient and r the remainder upon division of $\underline{x}$ by L.

Matching up the previous two equations, we see that if k-mer $\underline{x}$ is stored in its home slot q, then the value at address q is T[q] = L – r in the set {1,2,…,L}.

**2. Inserting and looking up keys**

Given a k-mer x and table T, first hash x to obtain $\underline{x}$, then compute the home slot q and remainder r, and then find the nearest available slot, scanning slot addresses sequentially upwards starting from the home address q. That is, find the smallest integer h (≥ 0) such that either $\underline{x}_{q+h} == \underline{x}$ or slot q + h is empty. In the first case, $\underline{x}$ is already in the table and there is nothing to do (unless we are also tracking the count of each k-mer; see [section] below). In the second case, if h ≤ H (the maximum allowed displacement from the home address), then write T[q+h] ← h * L – r to address q + h. If h > H, then signal that an overflow occurred, and write x to overflow table $T_{ov}$.

Pseudocode for inserting an element:

**Inputs**: hash table T, k-mer x

**Outputs**: 1) address at which x was inserted into T, 2) a Boolean signal denoting whether overflow occurred (i.e. too many hash collisions occurred)

```
def insertkey(T, x):
    x ← U * x (mod |X|^k)          # hash x
    q ← div(x, L)                  # quotient upon division by L
    r ← rem(x, L)                  # remainder upon division by L
    for h in 1 to H:               # at most H hash collisions
        if T[q] == 0:              # if slot is empty
            T[q] ← h * L - r       # write key to table
            return (q, false)      # no overflow
        else:                      # slot is occupied
            y ← (q + 1) * L - T[q] # hashed k-mer stored in slot
            if y == x:             # key is already in the hash table
                return (q, false)  # no overflow
        q ← q + 1                  # move to next slot
    return (q, true)               # overflow occurred
```

If an overflow occurs, then x is stored in an overflow table.

#### 3. Reading an element from the table

**Inputs**: hash table T, k-mer x

**Outputs**: 1) address at which x is in T, 2) a Boolean signal denoting whether x is in T (output 1 is undefined if this signal is false), 3) a signal denoting whether overflow occurred (i.e. too many hash collisions occurred)

```
def locatekey(T, x):
    x ← U * x (mod |X|^k)               # hash x
    q ← div(x, L)                       # quotient upon division by L
```

```
r ← rem(x, L)                       # remainder upon division by L
for h in 1 to H:                    # at most H hash collisions
    if T[q] == 0:                   # if slot is empty
        return (q, false, false)    # x not in table
    else:                           # slot is occupied
        y ← (q + 1) * L - T[q]      # hashed k-mer stored in slot
        if y == x:                  # key is already in the hash table
            return (q, true, false) # no overflow
     q ← q + 1                      # move to next slot
return (q, false, true)             # overflow occurred
```

If an overflow occurs, then check the overflow table for x.
